# Immunohistochemistry-based taxonomical classification of bladder cancer predicts response to neoadjuvant chemotherapy

**DOI:** 10.1101/724351

**Authors:** A. Font, M. Domènech, R. Benítez, M. Rava, M. Marqués, J. L. Ramírez, S. Pineda, S. Domínguez, J. L. Gago, J. Badal, C. Carrato, H. López, A. Quer, D. Castellano, N. Malats, F.X Real

**Affiliations:** Medical Oncology Service, B-ARGO Group, Institut Català d’Oncologia, Badalona, Spain; Medical Oncology Department, Fundació Althaia, Manresa, Spain; Genetic and Molecular Epidemiology Group, Spanish National Cancer Research Centre (CNIO), Madrid, Spain; CIBERONC, Spain; Epithelial Carcinogenesis Group, Spanish National Cancer Research Centre (CNIO), Madrid, Spain; IGTP-Molecular Biology Laboratory. Institute Germans Trias i Pujol, Badalona, Barcelona, Spain; Urology Department, Hospital Germans Trias i Pujol, Badalona, Spain; Pathology Department, Fundació Althaia, Manresa, Spain; Pathology Department, Hospital Germans Trias i Pujol, Badalona, Spain; Universitat Autònoma de Barcelona, Spain; Urology Department, Fundació Althaia, Manresa, Spain; Servicio de Oncología Médica, Hospital 12 de Octubre, Madrid, Spain; Departament de Ciències Experimentals i de la Salut, Universitat Pompeu Fabra, Barcelona, Spain

**Keywords:** bladder cancer, neoadjuvant chemotherapy, molecular taxonomy, immunohistochemistry, basal/squamous-like tumors

## Abstract

Platinum-based neoadjuvant chemotherapy (NAC) increases the survival of patients with organ-confined urothelial bladder cancer (UBC). Because not all patients benefit from treatment, NAC has not been widely applied in the clinical setting. There is strong evidence, based on retrospective studies, that patients with Basal/Squamous (BASQ)-like tumours present with more advanced disease and have worse prognosis; global transcriptomics can identify tumour subtypes associated with response to NAC. We aimed to investigate whether tumour immunohistochemical (IHC) subtyping predicts NAC response. Patients with muscle-invasive UBC having received platinum-based NAC were identified in two hospitals in Spain. Tissue microarrays were constructed; RNA and DNA were extracted from full sections. Nanostring analysis and immunohistochemistry to identify BASQ-like tumours and mutational analysis of UBC oncogenes. We used hierarchical clustering to classify 126 tumours and adjusted logistic regression to assess association with treatment response. Outcomes were progression-free survival and disease-specific survival; univariable and multivariate Cox regression models were applied. We found very high concordance between mRNA and protein for the 4 markers analyzed. We identified three main subgroups: BASQ-like (FOXA1/GATA3 low; KRT5/6/14 high), Luminal-like (FOXA1/GATA3 high; KRT5/6/14 low), and mixed-cluster (FOXA1/GATA3 high; KRT5/6 high; KRT14 low). Patients with BASQ-like tumours were more likely to achieve a pathological response to NAC, displaying a disease-specific survival similar to that of the remaining patients. In conclusion, patients with BASQ-like tumours - identified through simple and robust immunohistochemistry - have a higher likelihood of undergoing a pathological complete response to NAC. Prospective validation in independent series is required.

**Novelty and impact:** Neoadjuvant chemotherapy is an important component of the management of patietns with muscle-invasive bladder cancer but improved stratification is necessary. This retrospective study shows that patients with BASQ-like tumors can be identified using immunohistochemistry on paraffin-embedded tissue and are 4-fold more likely to achieve a pathological complete response to platinum-based NAC. The disease-specific survival of patients with BASQ-like tumours treated with NAC was not different from that of other tumour subtypes.

## Introduction

Cisplatin-based neoadjuvant chemotherapy (NAC) is the recommended treatment for locally-advanced muscle-invasive bladder cancer (MIBC), with highest level of evidence (1). Meta-analyses found that the benefit of NAC is limited to a subset of patients, with an improvement of 5-6.5% in 5-year survival compared to cystectomy alone (2,3). A recent meta-analysis shows a significant overall survival (OS) benefit associated with cisplatin-based NAC [hazard ratio (HR), 0.87; 95% confidence interval (CI), 0.79– 0.96] (4). Patients benefitting most from NAC are those who achieve a pathological complete response (pCR) at cystectomy (4,5). pCR or down-staging to non-MIBC (pT1-pTis) occur in almost half of patients treated with NAC. The inability to select patients who benefit has limited the use of NAC: treatment can lead to unnecessary toxicity in patients who fail to respond and it delays a potentially curative cystectomy, with a negative survival impact (6,7).

A better understanding of the molecular heterogeneity of UBC (8–10) has enabled classifying MIBC patients according to molecular profiles. Several groups have proposed classifications of all (11), NMIBC (12) or MIBC (8–10,13,14) tumours according to their transcriptome. Common to them is the identification of a tumour subgroup resembling basal-like breast cancers; there is consensus that Basal/Squamous-like (BASQ) tumours can be defined by ¡ KRT5/6 and KRT14 expression and lack of GATA3 and FOXA1 (15). Patients with BASQ-type UBC have more advanced disease at presentation and worse prognosis (13,16,17) but may respond better to NAC (13,17,18). The BASQ subtype was originally reported using global transcriptomics but its phenotype can be recognized with high accuracy using immunohistochemistry (IHC) (19,20). Whether adding urothelial differentiation markers could improve prediction is unknown.

Here, we assess the relationship between MIBC molecular IHC subtypes and response to NAC and prognosis in a retrospective series of patients treated with NAC.

## Materials and methods

### Study design

This study is based on a retrospective series of patients with MIBC treated with platinum-based NAC. The study included patients treated at two Spanish hospitals. MIBC was identified by transurethral resection of the bladder tumour (TURBT). Patients clinically staged with abdominal/pelvic computed tomography (CT) and a chest X-ray, and classified as T2-4aN0-2M0, were candidates for cystectomy after NAC. pCR was defined as absence of detectable tumour in the cystectomy specimen (pT0N0); partial response was defined as down-staging to non-MIBC (<pT2N0). Remaining cases were considered nonresponders. Patients provided informed consent; the study was approved by institutional review boards.

NAC was administered to 215 patients (1994-2014). The chemotherapy regimen was cisplatin, methotrexate and vinblastine (CMV) until 2000, after which cisplatin+gemcitabine (CG) or – exceptionally - carboplatin+gemcitabine (CaG) was used. 126 patients were eligible; remaining patients were excluded because tumour tissue was unavailable or insufficient for analyses. The main reason for lack of tissue availability was that one hospital is a referral centre and the TURBT had been performed elsewhere. Patients not undergoing surgery were excluded. Information was collected through retrospective review of clinical and pathological records.

### Immunohistochemiscal analyses

TURBT formalin-fixed paraffin-embedded (FFPE) blocks were reviewed. Representative tumour areas were selected to extract 3 cores and construct tissue microarrays (TMA). Cores were positioned non-consecutively to avoid artefacts; TMAs were constructed following established guidelines and sectioned; slides were embedded in paraffin and stored. The following antibodies were used: KRT5/6 (PRB-160P, Covance; 1/2000), KRT14 (PRB-155P, Covance, 1/2000), GATA3 (CM405 A, Biocare Medical, 1/300), FOXA1 (ab170933, Abcam, 1/100), FGFR3 (B9, Santa Cruz), KRT20 (Ks20.8, Dako), and STAG2 (J-12, Santa Cruz). After deparaffinization, antigen retrieval, and endogenous peroxidase blockade, sections were incubated with antibodies overnight, washed, and Envision secondary reagents (Agilent, Santa Clara, CA) was added for 1 h. After washing, reactions were developed with DAB, counterstained with haematoxylin, and mounted. Scoring was blind to clinical/pathological information. The proportion of reactive cells (0-100%) and staining intensity (0-3) were recorded; histoscore (HS) was calculated as the product (Intensity x % reactive cells, range 0-300). The average of the scores was used for analysis.

### RNA expression analyses

Total RNA was isolated from full FFPE sections and purified from macrodissected high-density tumour areas using the truxTrac FFPE RNA microTube kit (Covaris). Cell lysates were sheared by sonication; RNA was eluted and quantified using Qubit. Gene expression analysis was conducted on the NanoString nCounter platform. Samples were scanned at maximum resolution on the nCounter Digital Analyzer. Four housekeeping transcripts were used for normalization (*GAPDH*, *ACTB*, *HPRT1*, and *LDHA*).

### Mutational analyses

Hotspot mutations in *FGFR3*, *PIK3CA*, and *HRAS*, *KRAS* and *NRAS* were analyzed. DNA was extracted from tumour cell-enriched areas using DNeasy tissue kit (Qiagen, Hilden, Germany). Multiplex PCRs were performed; products were analysed with the SNaPshot Multiplex Kit (Applied Biosystems, Foster City, CA) (21).

### Statistical methods

Variables were summarized as means, medians and standard deviations, when continuous, and as percentages when categorical. Bivariable associations were evaluated by chi2 test (categorical), and Kruskal-Wallis test (continuous). Agglomerative hierarchical clustering was applied using the Ward’s minimum variance method (Ward 1963) to investigate evidence for natural groupings of tumours (subtypes) based on correlations between expression profiles. HS were standardized to a 0-1 and considered as input. Heatmaps were generated using pHeatmap (22). To investigate cluster stability, bootstrapped HS values were re-clustered and their similarity with the original classification was evaluated with the Jaccard coefficient (23). CART model was trained considering the 3 subphenotypes and the 4 markers using caret and rpart packages. *Leave-one-out cross validation* was used to evaluate the best prediction scenario.

Adjusted logistic regression was applied to assess the independent association between clusters and response. For each variable, the odds ratio (OR) was estimated referring to the association between the predictor pCR with the ‘no-complete response’ group - including partial responders and non-responders - as reference. Sensitivity analyses were run after excluding subjects with pelvic lymph node involvement or treated with carboplatin (N=31).

Mean follow-up in event-free patients was 85 months (6 - 241). Fifty patients (37.9%) underwent progression and 45 (34.1%) died from disease. Survival curves were derived using Kaplan-Meier methods and compared with log-rank tests by strata defined by tumour subtypes obtained with cluster analysis. Disease specific-survival and progression-free survival were considered as outcomes. Hazard ratios and 95% confidence interval for the association between subtypes and outcomes were estimated with univariable and multivariable Cox regression models.

Differences were considered statistically significant at P<0.05. All statistical analyses were performed using R version 3.3 unless other otherwise indicated.

## Results

### Clinical-pathological features of patients

Clinical/pathological characteristics of patients included in the study (n=132) and those from whom no material was available (n=83) were similar, indicating that the cases analysed are representative of the whole population. The only significant difference was the NAC regimen used, reflecting variability related to the inclusion period (Supplementary Table 1). Six tumours were excluded due to lack of information on all markers. Baseline characteristics of 126 patients with IHC data are listed in Table 1. Median age was 66 (range, 61-72); 86% were pure urothelial carcinomas; the remaining 14% had mixed histology, with squamous predominance. Clinical staging was T2-4N0M0 for 109 (83%) patients; 21 (17%) had nodal involvement. Chemotherapy regimens used were: CG (63%), CMV (25%), and CaG (10%).

**Table 1.**
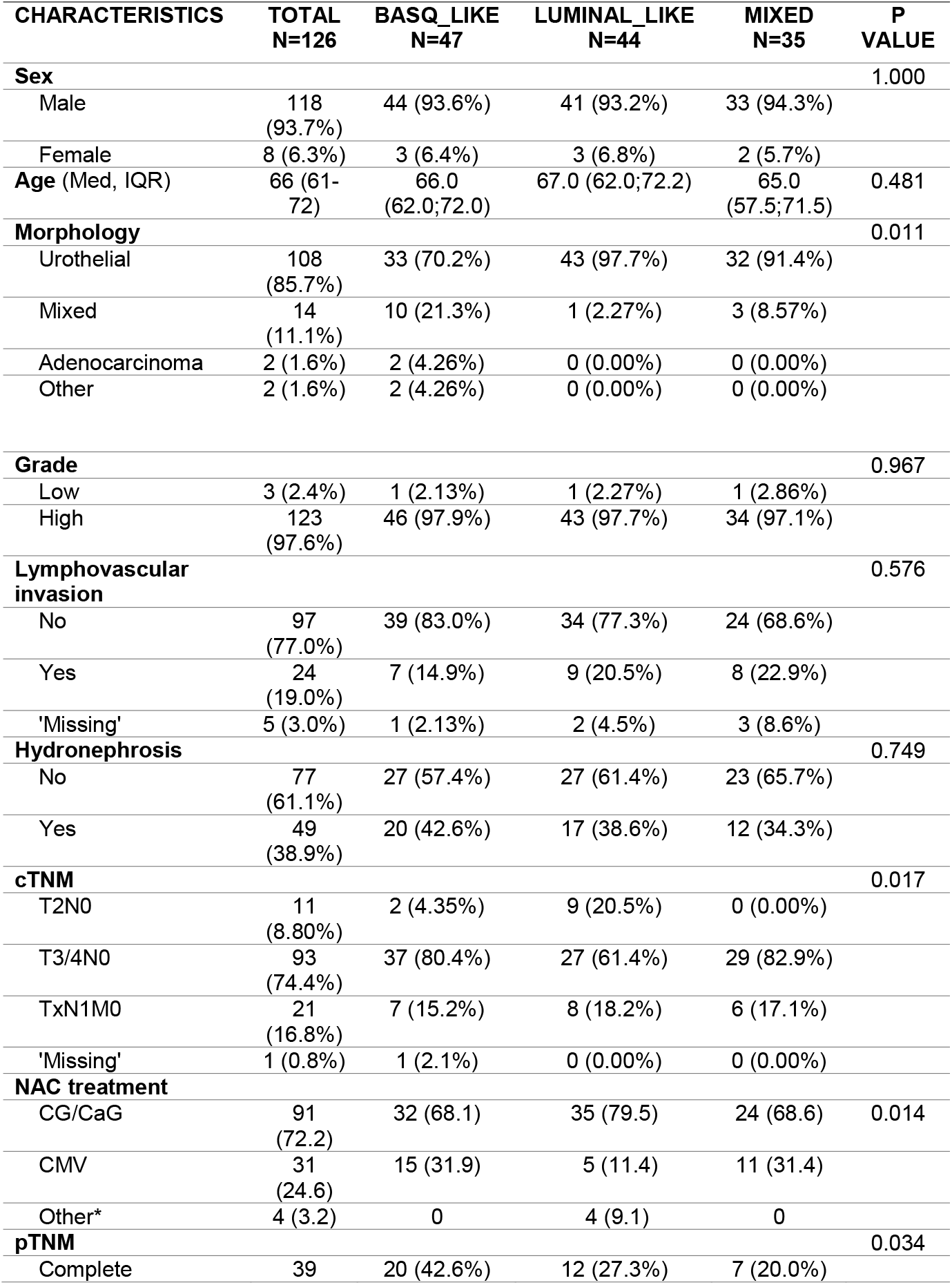
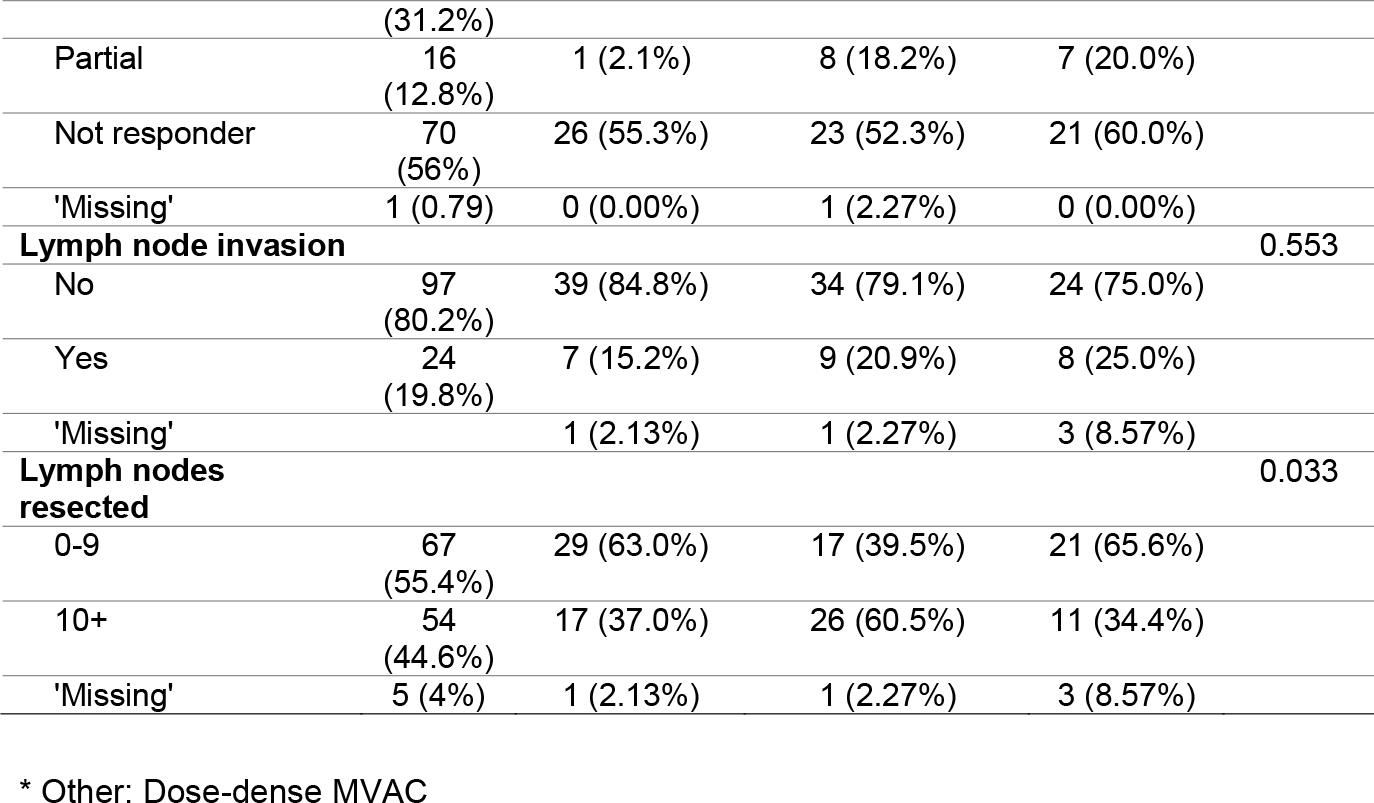
Patient and tumour characteristics according to the taxonomic clusters (N=126).

Pathologic down-staging occurred in 57 (43%) patients; 40 cases (30%) had a pCR. Baseline characteristics of responders and non-responders were similar except for nodal involvement, which was more common among non-responders (26% vs 0%; P=0.002) (Supplementary Table 2).

### Taxonomical classification and association with gene mutations

We compared Nanostring RNA quantification of the four markers with protein expression using IHC. Highly significant Spearman correlation coefficients were found for GATA3, FOXA1, and KRT14 and good correlation for KRT5/6 (Figure 1A).

**Figure 1.**
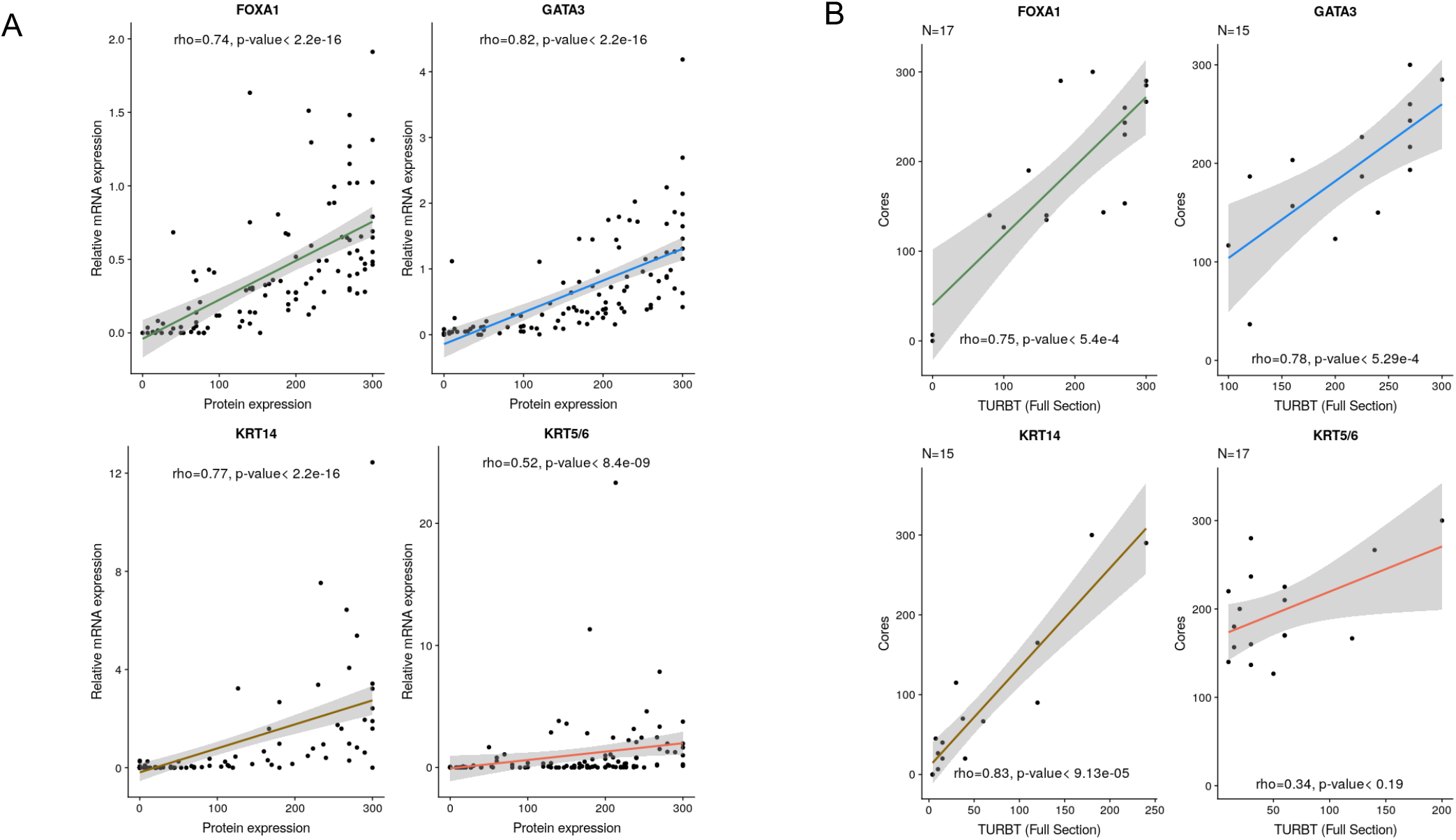
**(A)** Comparison of the results of IHC findings and Nanostring expression for the markers used for tumour subtype classification shows excellent correlation between both assays. **(B)** Comparison of the HS of TMA cores with the findings in full sections of the corresponding TURBT full block sections.

IHC expression of FOXA1 and GATA3 showed a significant positive correlation (Pearson r=0.68). KRT5/6 and KRT14 expression were also significantly positively correlated (r=0.45). FOXA1 and GATA3 were negatively correlated with KRT14 (r=-0.53 and r= −0.62, respectively). GATA3, but not FOXA1, expression correlated negatively with KRT5/6 (r=-0.33). Intercore HS concordance analysis revealed moderate (0.4-0.6) or substantial (>0.6) agreement for all markers. Concordance was higher for the BASQ group than for the luminal or mixed groups.

To avoid the need of using thresholds for subtype classification, hierarchical clustering was used and identified three clusters. Cluster 1 (BASQ-like) (N=47) is characterized by high expression of KRT5/6 and KRT14 and low expression of FOXA1 and GATA3. Cluster 2 (Urothelial/Luminal-like) (N=44) is characterized by high expression of FOXA1 and GATA3 and low expression of KRT5/6 and KRT14. Cluster 3 (mixed-cluster) (N=35) is characterized by high expression of FOXA1, GATA3 and KRT5/6, and low expression of KRT14 (Figure 2A). Re-clustering of bootstrapped protein-expression showed that the “BASQ-like” cluster was the most stable (stability = 0.84). CART analysis showed that KRT14 and KRT5/6 expression provided the best cluster discrimination (Figure 2B). In agreement with global transcriptome analyses, a few tumours were identified with low expression of all markers but they did not form a distinct cluster.

**Figure 2.**
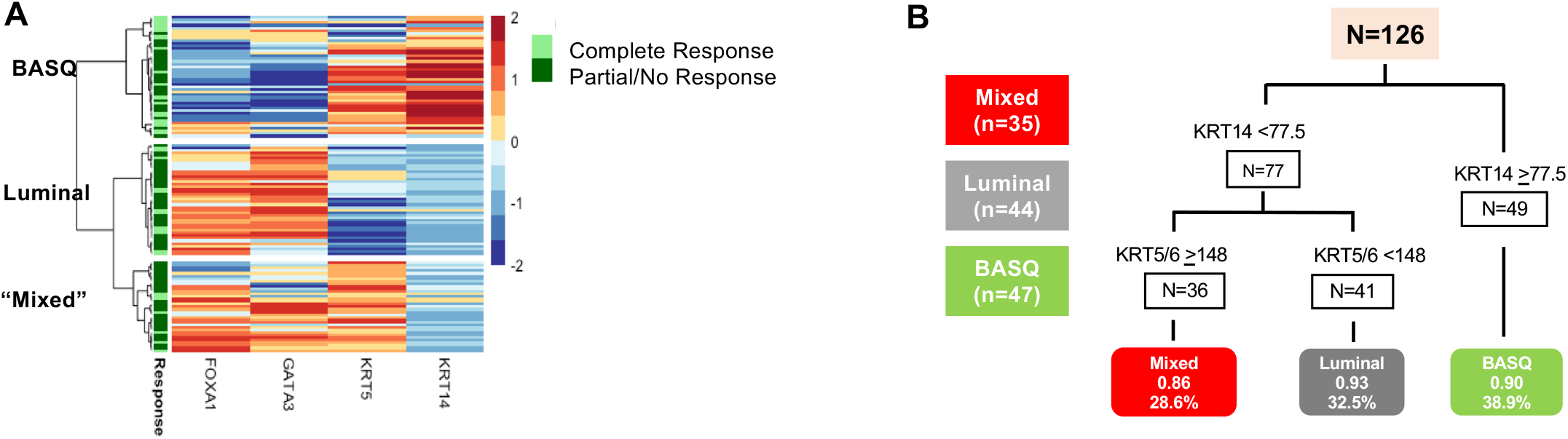
Expression of FOXA1, GATA3, KRT5/6 and KRT14 in the three tumour subtypes. The heatmap depicts relative biomarker expression. **(A)** Clusters resulting from KRT5/6, KRT14, GATA3, and FOXA1 tumour HS. **(B)** CART model showing the optimal accuracy of cluster classification.

BASQ-like tumours were enriched for subjects with more advanced clinical stage at diagnosis (P=0.013), mixed squamous histology (P=0.011), higher number of resected nodes (P=0.033), and higher pCR rate (P=0.034).

The first molecular taxonomy consensus classification did not include markers of luminal/urothelial differentiation (15). Among the candidate molecules associated with urothelial differentiation are KRT20, a well-established luminal marker and *FGFR3, STAG2*, and *PIK3CA* - three major bladder cancer genes commonly altered in luminal tumors (21,24). Therefore, we analyzed these markers in the same samples. KRT20 expression was significantly lower in BASQ tumours (Supplementary Table 3) but it was similar in the other clusters. FGFR3 expression was significantly higher in the “mixed-cluster” tumours and STAG2 expression was similar in all clusters. *FGFR3* and *PIK3CA* mutations were more common in the mixed-cluster tumours; differences did not reach statistical significance. *RAS* mutations were similarly distributed across clusters (Supplementary Table 4).

### Association of clusters with response to NAC

To analyse the association with response to NAC, we focused on the consensus markers of the BASQ-like cluster. Supplementary Table 2 shows the clinical variables and molecular factors associated with pCR in the univariable analysis. Table 2 shows factors predictive of pCR in the multivariate analysis. The two models differ in the inclusion of lymphovascular invasion and lymph node metastases. Young age and BASQ-like cluster were the main variables positively associated with pCR. In both models, the OR for BASQ-like tumours was approximately 4.

**Table 2.** Predictive factors of complete response to neoadjuvant treatment. Multivariate logistic regression models (N=126)

| Factors | Model 1 |  |  | Model 2 |  |  |
| --- | --- | --- | --- | --- | --- | --- |
|  | OR | P value | 95% CI | OR | P value | 95% CI |
| Morphology |  |  |  |  |  |  |
| Urothelial | Ref |  |  | Ref |  |  |
| Mixed | 0.27 | 0.083 | (0.06,1.19) | 0.32 | 0.159 | (0.06,1.56) |
| Lymphovascular invasion |  |  |  |  |  |  |
| No | - |  |  | Ref. |  |  |
| Yes | - |  |  | 0.61 | 0.531 | (0.13,2.90) |
| Lymph node invasion |  |  |  |  |  |  |
| No | - |  |  | Ref |  |  |
| Yes | - |  |  | 0.07 | 0.014 | (0.00,Inf) |
| cTNM |  |  |  |  |  |  |
| T2 N0 | Ref |  |  | Ref |  |  |
| T3/4 N0 | 0.31 | 0.121 | (0.07,1.37) | 0.22 | 0.093 | (0.04,1.28) |
| TxN1Mx | 0.36 | 0.245 | (0.06,2.01) | 0.33 | 0.293 | (0.04,2.59) |
| Clusters |  |  |  |  |  |  |
| Mixed | Ref |  |  | Ref |  |  |
| Luminal-like cluster | 1.28 | 0.699 | (0.37,4.49) | 1.34 | 0.672 | (0.34,5.26) |
| BASQ-like cluster | **3.96** | **0.017** | (1.28,12.20) | **4.06** | **0.026** | (1.18,13.99) |

Both models were adjusted for age and centre.

A few patients included had lymph node involvement or received carboplatin, which are not part of standard NAC. When these cases were excluded (sensitivity analysis), the predictive value of the BASQ-like cluster was essentially unchanged (Supplementary Table 5).

Figure 3 shows Kaplan-Meier plots for relapse-free and disease-specific survival. Among all patients and among those who underwent a pCR, cluster classification was not associated with survival; patients with BASQ tumours having achieved a pCR had the best survival. By contrast, among patients who did not have a pCR, those with BASQ tumours had worst outcome (P=0.11). These findings strongly suggest that the worse prognosis of patients with BASQ-like tumours is counter-effected by a greater likelihood of displaying a pCR, resulting in lack of differences in disease-specific survival.

**Figure 3.**
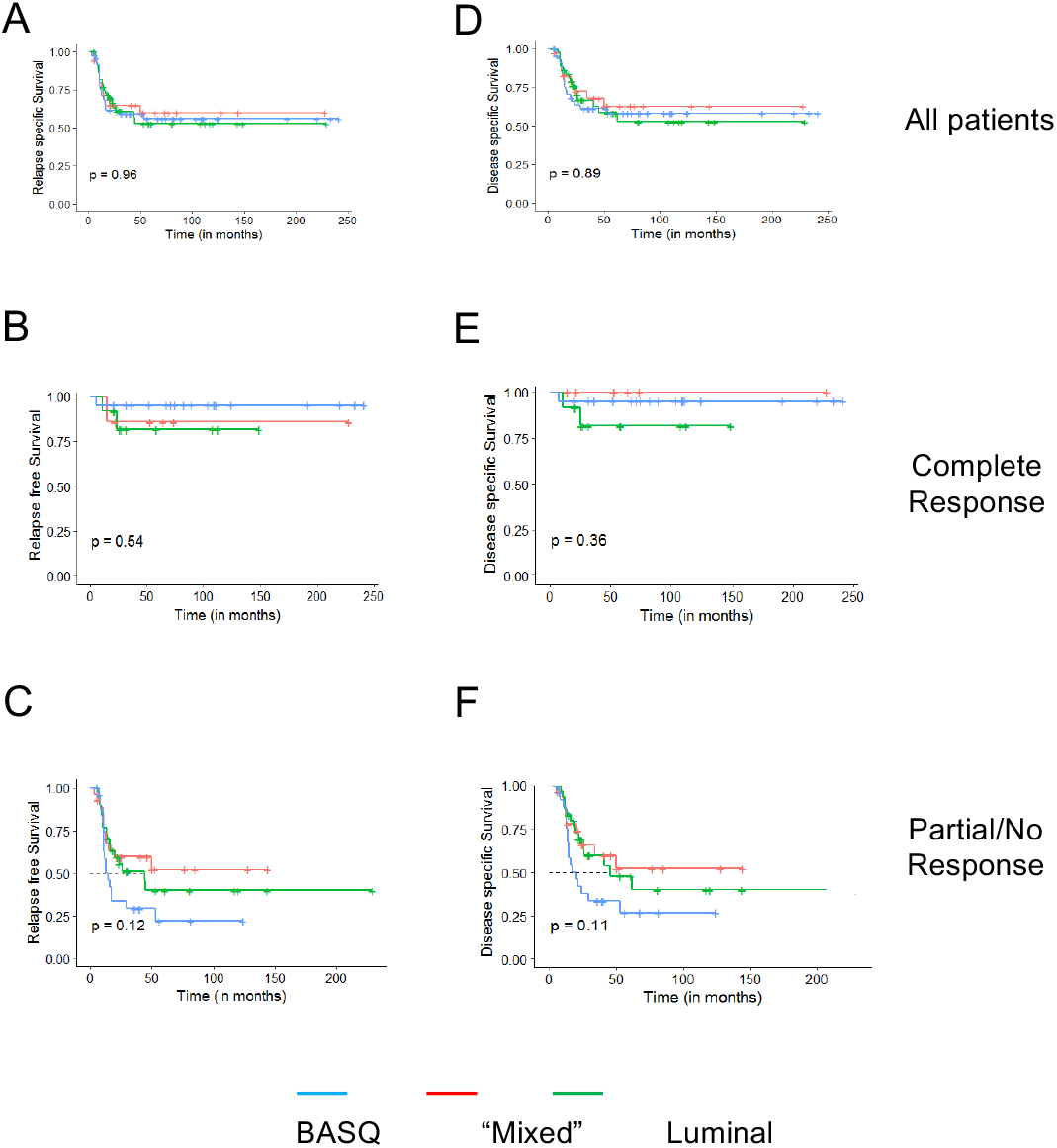
Kaplan-Meier disease-specific survival curves for all patients **(A)**, patients undergoing a pCR **(B)**, or partial/non-responders **(C)** according to the three tumour subtypes.

### Relevance to clinical implementation

Because clinical implementation would not rely on TMAs, we compared marker expression in TMA vs. full TURBT sections (n=15) and found highly significant correlations for FOXA1 (r = 0.75), GATA3 (r=0.78), and KRT14 (r=0.83) and a modest correlation for KRT5/6 (r=0.34) (Figure 1B). These correlations validate, overall, the usefulness of the TMAs for exploratory studies and agree with a recent publication (25). To determine whether TURBT are representative of the whole tumour, we compared marker expression in full sections of paired TURBT and cystectomy specimens from patients receiving NAC who did not achieve a pCR (n=17) and from patients who did not receive NAC (n=15). We find a significantly reduced HS for luminal markers and a consistent - less prominent - increased expression of KRT basal markers. These results suggest heterogeneity in the cellular phenotypes in superficial vs. deeper tumour regions (Supplementary Figures 1,2). Interestingly, the trend of the changes in KRT14 expression were different in samples that had been exposed, or not, to NAC suggesting an impact of the latter on tumour cell phenotype (Supplementary Figure 3).

## Discussion

Molecular knowledge has generated a plethora of hypotheses that could impact on UBC patient management. The fact that only a fraction of patients respond to available therapies, highlights the need of improved patient stratification - especially with the advent of immunotherapy. This need is obvious for NAC, where the risk of tumour progression in non-responders has underscored its cautious application even when global survival statistics supports its use in patients with locally advanced organ-confined tumours.

In MIBC, several tumour subtypes have been identified; greatest consensus exists for the BASQ-like type, defined using transcriptomics. Based on this, and on the increasing evidence that BASQ tumours are more aggressive but may respond better to chemotherapy, we performed this analysis using IHC.

To our knowledge, ours is the first study showing that a robust cluster of tumours with BASQ-like features, approximately one-third of all patients with MIBC, can be identified using simple IHC to predict NAC response. This cluster is defined by low/undetectable expression of GATA3/FOXA1 and high expression of KRT5/6/14. KRT5/6 did not clearly distinguish BASQ-like tumours from the mixed-cluster tumour subtype. These markers have also previously been shown by the Lund group to robustly translate global transcriptomic classifiers into immunohistochemical assays (19). Because of the well-defined specificity of the antibodies used and the robustness of the IHC stainings, the assay combination used here could be widely applied in Pathology Departments, facilitating clinical translation. Patients with BASQ-like tumours have a significantly higher likelihood of response to NAC in comparison to the other tumour subgroups, but only 42% of them underwent a pCR, indicating that further refinement is required to improve patient selection.

Three strategies have been applied to identify markers of response to cisplatin-based NAC: gene mutation (26,27), global transcriptomics (14,17), and panel IHC (here). Somatic *ERCC2* mutations are enriched in NAC-responders and functional *in vitro* studies supported a causal association (26). Genomic alterations in *RB1*, *ATM*, and *FANCC* were predictive of response to, and clinical benefit from, NAC (27).

Two studies used global transcriptomics using FFPE samples to define UBC subtypes in relationship to response to NAC. Patients with basal tumours from a clinical trial with dose-dense MVAC and bevacizumab had improved survival compared to those with luminal or p53-like tumours. In that study, NAC response was not associated with tumour subtype (18). In a large multicentre retrospective series comparing patients having undergone cystectomy without NAC and those having received NAC, patients with basal tumours not receiving NAC consistently had worse overall survival than those with luminal tumours. Importantly, patients with basal-like tumours (SCC-like, cluster III, or Uro B) showed greatest improvement in outcome after NAC. In multivariate analysis, patients with basal tumours not receiving NAC had an adjusted HR of 2.22 for OS compared to luminal tumours; these differences were eroded in the NAC cohort. Surprisingly, among patients with basal tumours (unlike in luminal tumours), OS was similar for major responders vs. non-responders (17). The use of group comparisons is complicated by the fact that some patients not undergoing NAC receive adjuvant chemotherapy or - eventually - receive cisplatin-based chemotherapy upon if they progress during follow-up.

Our study relied on the type of samples that would be used for patient stratification, i.e. TURBT. We find that these samples may not be completely representative of the whole tumour. The deep region thereof, sampled at cystectomy, tends to have a more basal phenotype, highlighting tumour heterogeneity. A recent study of bladder cancer histological variants has shown that BASQ tumors display greatest heterogeneity and supports that variants arise from conventional urothelial carcinomas (28). More work needs to be carried out to specifically assess the relevance of sample bias, the difference between primary tumour vs. lymph node or distant metastases (29,30) as well as the changes in tumor biology associated with tumor evolution, naturally or under therapy pressure (31). Furthermore, prospective studies should determine the optimal strategy for BASQ tumor definition, considering the more recently proposed classifiers (32).

An important question is whether gene mutations are related to tumour subtypes. *ERCC2* mutation prevalence is similar in BASQ-like and non-BASQ-like tumours (8). *RB1* mutations – but not deletions – are significantly more common in BASQ-like than in urothelial tumours. However, deletions are more common in the Lund “genomically unstable” subgroup. There is insufficient data to indicate whether *ATM* and *FANCC* alterations are enriched in any taxonomical types (8,10), suggesting that combined mutational analysis and tumour subtyping could increase the ability to predict NAC response.

This study has several limitations derived from its retrospective nature and not being a clinical trial, including the relatively long recruitment period and modest sample size, lack of centralized pathology review, inclusion of patients treated with carboplatin, and possible uncontrolled differences in clinical management. These limitations have been mitigated through several types of analyses and they reflect much of the variability associated with the application of NAC in clinical practice (33,34). While the P-values found are close to the threshold of significance, the OR are in the range of clinical relevance. Nevertheless, prospective clinical studies are necessary to conclusively establish the contribution of tumour subtyping to the management of patients with MIBC and the optimal clinical implementation strategy.

## Conclusions

This retrospective study of patients with muscle-invasive bladder cancer shows that BASQ-like tumours can be reliably identified using IHC on FFPE sections, allowing the definition of a patient subgroup that is almost 4-fold more likely to respond to platinum-based NAC. Our data supports the conduct of a prospective study to confirm that patients with BASQ-like tumours benefit most from platinum-based NAC.

## Author contributions

*Study concept and design*: Castellano, Doménech, Font, Malats, Real

*Acquisition of data*: Badal, Carrato, Doménech, Font, Gago, López, Malats, Marqués, Quer, Ramírez, Real

*Analysis and interpretation of data*: Doménech, Font, Malats, Pineda, Ramírez, Real

*Drafting of the manuscript*: Doménech, Font, Malats, Real

*Critical revision of the manuscript for important intellectual content*: All authors

*Statistical analysis*: Rava, Benítez, Pineda, Domínguez, Malats

*Obtaining funding*: Castellano, Font, Malats, Real

*Supervision*: Malats, Real

## Acknowledgements

We thank Tania Lobato, José M. Velarde, and Natalia del Pozo for valuable contributions.

## Data availability statement

All data from this study are available upon reasonable request to authors.

