## Supplementary Materials for "Immunohistochemistry-based taxonomical classification of bladder cancer predicts response to neoadjuvant chemotherapy"

**Font et al.**

### Supplementary material

**Supplementary Table 1. Characteristics of the subjects included and excluded from the analyses**

| Characteristics | All N=215 | Excluded N=83 | Included N=132 | P |
|---|---|---|---|---|
| **Centre** | | | | 0.00016 |
| Can Ruti | 168 (78.1%) | 76 (91.6%) | 92 (69.7%) | |
| Manresa | 47 (21.9%) | 7 (8.43%) | 40 (30.3%) | |
| **Sex** | | | | 0.33 |
| M | 199 (92.6%) | 75 (90.4%) | 124 (93.9%) | |
| F | 16 (7.44%) | 8 (9.64%) | 8 (6.06%) | |
| **Age** (cont) | 66.0 [59.0;71.5] | 65.0 [58.0;70.5] | 66.0 [61.0;72.0] | 0.34 |
| **Age** (tertiles, years) | | | | 0.25 |
| [35,62] | 76 (35.3%) | 35 (42.2%) | 41 (31.1%) | |
| (62,69] | 72 (33.5%) | 25 (30.1%) | 47 (35.6%) | |
| (69,83] | 67 (31.2%) | 23 (27.7%) | 44 (33.3%) | |
| **Morphology** | | | | 0.37 |
| Urothelial | 183 (85.1%) | 70 (84.3%) | 113 (85.6%) | |
| Mixed | 23 (10.7%) | 9 (10.8%) | 14 (10.6%) | |
| Adenocarcinoma | 6 (2.79%) | 4 (4.82%) | 2 (1.52%) | |
| Other | 3 (1.40%) | 0 (0.00%) | 3 (2.27%) | |
| **Lymphovascular invasion** | | | | 0.261 |
| No | 188 (87.4%) | 74 (89.2%) | 114 (86.4%) | |
| Yes | 24 (11.2%) | 7 (8.43%) | 17 (12.9%) | |
| Unknown | 1 (0.47%) | 1 (1.20%) | 0 (0.00%) | |
| ‘Missing’ | 2 (0.93%) | 1 (1.20%) | 1 (0.76%) | |
| **Grade** | | | | 0.55 |
| Low | 6 (2.79%) | 3 (3.61%) | 3 (2.27%) | |
| High | 208 (96.7%) | 79 (95.2%) | 129 (97.7%) | |
| 'Missing' | 1 (0.47%) | 1 (1.20%) | 0 (0.00%) | |
| **cTNM** | | | | |
| 0-3 | 16 (7.44%) | 3 (3.61%) | 13 (9.85%) | 0.16 |
| 4-6 | 157 (73.0%) | 61 (73.5%) | 96 (72.7%) | |
| 7+ | 41 (19.1%) | 19 (22.9%) | 22 (16.7%) | |
| 'Missing' | 1 (0.47%) | 0 (0.00%) | 1 (0.76%) | |
| **pTNM**: | | | | 0.73 |
| Not assessable | 9 (4.19%) | 8 (9.64%) | 1 (0.76%) | |
| Complete | 58 (27.0%) | 18 (21.7%) | 40 (30.3%) | |
| Partial | 27 (12.6%) | 10 (12.0%) | 17 (12.9%) | |
| Not responder | 117 (54.4%) | 43 (51.8%) | 74 (56.1%) | |
| 'Missing' | 4 (1.86%) | 4 (4.82%) | 0 (0.00%) | |
| **Lymph node involvement** | | | | 0.04 |
| No | 161 (74.9%) | 59 (71.1%) | 102 (77.3%) | |
| Yes | 29 (13.5%) | 5 (6.02%) | 24 (18.2%) | |
| 'Missing' | 25 (11.6%) | 19 (22.9%) | 6 (4.55%) | |
| **Lymph nodes resected** | | | | 0.18 |
| 0-9 | 112 (52.1%) | 42 (50.6%) | 70 (53.0%) | |
| 10+ | 78 (36.3%) | 22 (26.5%) | 56 (42.4%) | |
| 'Missing' | 25 (11.6%) | 19 (22.9%) | 6 (4.55%) | |

| Characteristics | All<br>N=215 | Excluded<br>N=83 | Included<br>N=132 | P |
|---|---|---|---|---|
| **Treatment** | | | | 0.04 |
| CMV | 66 (30.7%) | 34 (41.0%) | 32 (24.2%) | |
| CG | 122 (56.7%) | 38 (45.8%) | 84 (63.6%) | |
| CaG | 20 (9.30%) | 8 (9.64%) | 12 (9.09%) | |
| Other* | 5 (2.33%) | 1 (1.20%) | 4 (3.03%) | |
| 'Missing' | 2 (0.93%) | 2 (2.41%) | 0 (0.00%) | |

* Other: Dose-dense MVAC

**Supplementary Table 2. Patient characteristics according to the response to treatment (N=131)***

| Characteristics | Complete N=40 | Partial/ Non-responders N=91 | P value |
|---|---|---|---|
| **Centre** | | | |
| Can Ruti | 28 (70.0%) | 64 (70.3%) | 1.000 |
| Manresa | 12 (30.0%) | 27 (29.7%) | |
| **Sex** | | | |
| M | 39 (97.5%) | 84 (92.3%) | 0.442 |
| F | 1 (2.50%) | 7 (7.69%) | |
| **Age** (cont.), median [1st-3rd quartile] | 66.0 [62.8;69.8] | 67.0 [60.5;72.0] | 0.777 |
| **Age** (tertiles, years) | | | |
| [35,62] | 10 (25.0%) | 31 (34.1%) | 0.067 |
| (62,69] | 20 (50.0%) | 26 (28.6%) | |
| (69,83] | 10 (25.0%) | 34 (37.4%) | |
| **Morphology** | | | |
| Urothelial | 37 (92.5%) | 75 (82.4%) | 0.465 |
| Mixed | 3 (7.50%) | 11 (12.1%) | |
| Adenocarcinoma | 0 (0.00%) | 2 (2.20%) | |
| Other | 0 (0.00%) | 3 (3.30%) | |
| **Lymphovascular invasion** | | | |
| No | 36 (90.0%) | 77 (84.6%) | 0.568 |
| Yes | 4 (10.0%) | 13 (14.3%) | |
| Unknown | 0 (0.00%) | 1 (1.10%) | |
| **Grade** | | | |
| Low | 1 (2.50%) | 2 (2.20%) | 1.000 |
| High | 39 (97.5%) | 89 (97.8%) | |
| **cTNM** | | | |
| 0-3 | 6 (15.0%) | 6 (6.59%) | 0.303 |
| 4-6 | 27 (67.5%) | 69 (75.8%) | |
| 7+ | 6 (15.0%) | 16 (17.6%) | |
| 'Missing' | 1 (2.50%) | 0 (0.00%) | |
| **Lymph node involvement** | | | |
| No | 38 (95.0%) | 63 (69.2%) | 0.002 |
| Yes | 0 (0.00%) | 24 (26.4%) | |
| ‘Missing’ | 2 (5.00%) | 4 (4.40%) | |
| **Lymph nodes resected** | | | |
| 0-9 | 19 (47.5%) | 51 (56.0%) | 0.449 |
| 10+ | 19 (47.5%) | 36 (39.6%) | |
| 'Missing' | 2 (5.00%) | 4 (4.40%) | |
| **Hydronephrosis** | | | |
| No | 27 (67.5%) | 52 (57.1%) | 0.339 |
| Yes | 13 (32.5%) | 39 (42.9%) | |
| **Treatment** | | | |
| CG | 25 (62.5%) | 58 (63.7%) | 1.000 |
| CaG | 4 (10.0%) | 8 (8.79%) | |
| CMV | 10 (25.0%) | 22 (24.2%) | |
| 'Missing' | 1 (2.50%) | 3 (3.30%) | |

*For one patient, response could not be assessed.

**Supplementary Table 3. Median and interquartile range [IQR] of HS corresponding to each tumour marker according to the 3 clusters identified (N=126)**

| Marker | **Mixed** N=35 | **Luminal-like** N=44 | **BASQ-like** N=47 | **P value** |
|---|---|---|---|---|
| FOXA1 | 220 [145;288] | 250 [187;272] | 80.0 [40.0;147] | 1.71E-11 |
| GATA3 | 193 [155;242] | 265 [219;290] | 53.3 [15.0;153] | 3.32E-13 |
| KRT5 | 220 [195;240] | 77.5 [20.0;122] | 207 [135;250] | 9.20E-14 |
| KRT14 | 20.0 [0.00;47.5] | 0.00 [0.00;20.0] | 217 [152;275] | 2.05E-17 |
| KRT20 | 93.3 [0.00;155] | 71.7 [18.8;140] | 0.00 [0.00;63.3] | 0.001 |
| FGFR3 | 33.3 [10.0;90.0] | 10.0 [0.00;38.3] | 6.67 [0.00;37.5] | 0.021 |
| STAG2 | 233 [197;273] | 267 [238;300] | 250 [170;300] | 0.242 |

**Supplementary Table 4. Distribution of bladder cancer gene mutations according to the 3 clusters identified (N=126)**

| MARKER | MIXED N=35 | LUMINAL-LIKE N=44 | BASAL-LIKE N=47 | P VALUE |
|---|---|---|---|---|
| *FGFR3* | | | | 0.015 |
| wt | 23 (69.7%) | 35 (79.5%) | 44 (93.6%) | |
| mut | 8 (24.2%) | 4 (9.09%) | 1 (2.13%) | |
| 'Missing' | 2 (6.06%) | 5 (11.4%) | 2 (4.26%) | |
| *PIK3CA* | | | | 0.132 |
| wt | 21 (63.6%) | 35 (79.5%) | 35 (77.8%) | |
| mut | 7 (21.2%) | 3 (6.82%) | 2 (4.44%) | |
| 'Missing' | 5 (15.2%) | 6 (13.6%) | 8 (17.8%) | |
| *HRAS* | | | | 0.314 |
| wt | 13 (39.4%) | 23 (54.8%) | 19 (42.2%) | |
| mut | 1 (3.03%) | 2 (4.76%) | 0 (0.00%) | |
| 'Missing' | 19 (57.6%) | 17 (40.5%) | 26 (57.8%) | |
| *KRAS* | | | | 0.131 |
| wt | 14 (42.4%) | 25 (58.1%) | 16 (35.6%) | |
| mut | 0 (0.00%) | 1 (2.33%) | 3 (6.67%) | |
| 'Missing' | 19 (57.6%) | 17 (39.5%) | 26 (57.8%) | |
| *NRAS* | | | | 0.241 |
| wt | 13 (39.4%) | 26 (60.5%) | 18 (40.0%) | |
| mut | 1 (3.03%) | 0 (0.00%) | 1 (2.22%) | |
| 'Missing' | 19 (57.6%) | 17 (39.5%) | 26 (57.8%) | |
| *RAS*-any | | | | 0.415 |
| wt | 12 (36.4%) | 22 (52.4%) | 15 (33.3%) | |
| mut | 2 (6.06%) | 3 (7.14%) | 4 (8.89%) | |
| 'Missing' | 19 (57.6%) | 17 (40.5%) | 26 (57.8%) | |

**Supplementary Table 5. Predictive factors of pathological complete response to neoadjuvant treatment. Multivariate logistic regression models (N=100*)**

| Factor | Model 1 | | Model 2 | |
|---|---|---|---|---|
| | **OR** | **P value** | **OR** | **P value** |
| Age (years) | | | | |
| <62 | Ref | | Ref | |
| 62-69 | 2.10 | 0.186 | 2.16 | 0.206 |
| 70-83 | 0.69 | 0.560 | 0.76 | 0.693 |
| Morphology | | | | |
| Urothelial | Ref | | Ref | |
| Mixed | 0.43 | 0.310 | 0.54 | 0.471 |
| Lymphovascular invasion | | | | |
| No | – | – | Ref | |
| Yes | – | – | 1.38 | 0.723 |
| Lymph node involvement | | | | |
| No | – | – | Ref | |
| Yes | – | – | 0.08 | 0.03 |
| cTNM | | | | |
| T2 N0 | Ref | | Ref | |
| T3/4 N0 | 0.33 | 0.190 | 0.19 | 0.093 |
| Clusters | | | | |
| Mixed | Ref | | Ref | |
| Luminal-like cluster | 0.93 | 0.916 | 0.81 | 0.790 |
| BASQ-like cluster | 3.22 | 0.062 | 3.28 | 0.076 |

* Patients treated with Carboplatin or with lymph node involvement were excluded from the analyses. Both models were adjusted for centre.

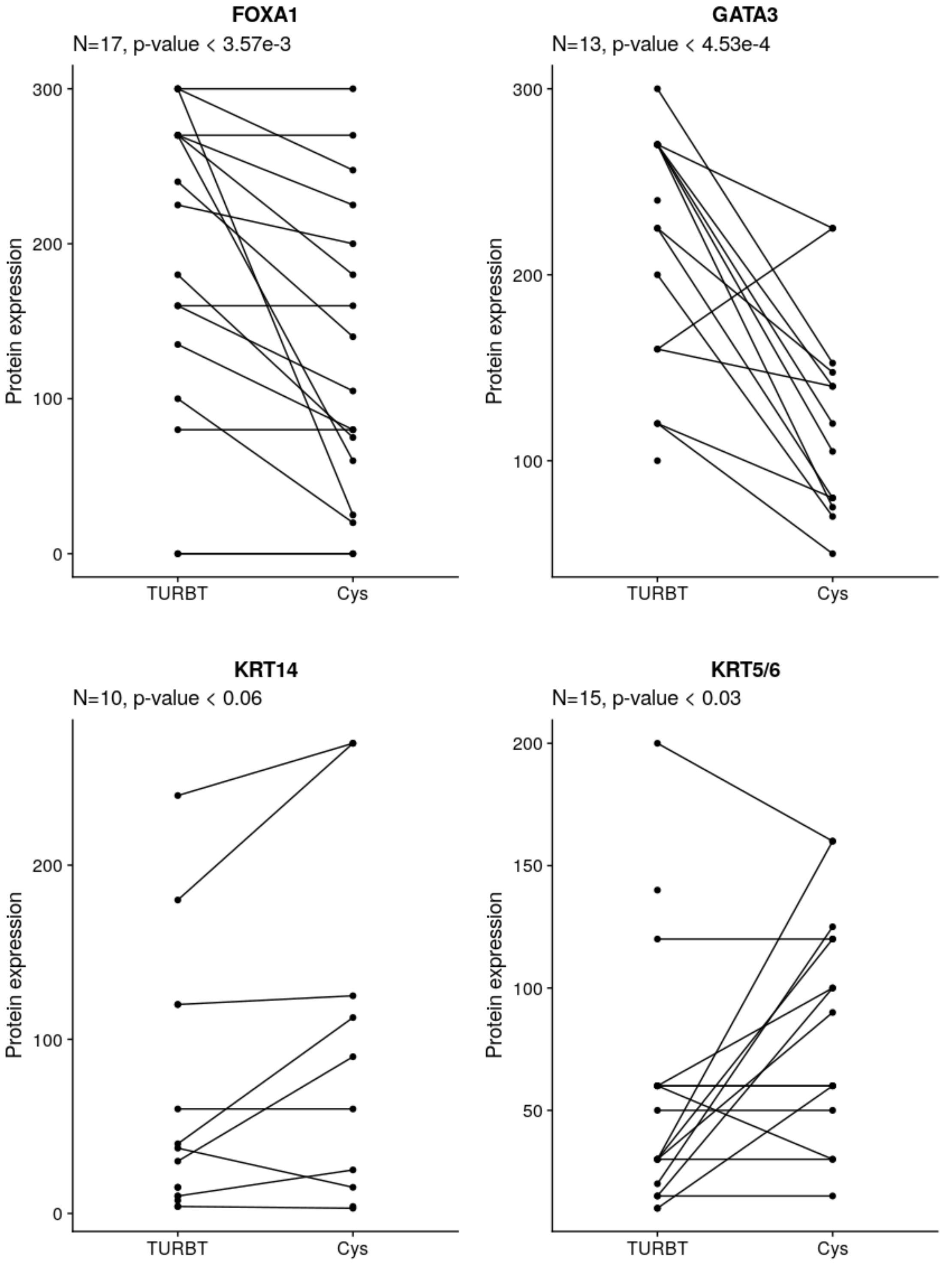

**Supplementary Figure 1**. Comparison of marker expression in full sections of samples obtained at TURBT and cystectomy (Cys) from patients treated with NAC. Statistical analysis: repeated measurements ANOVA.

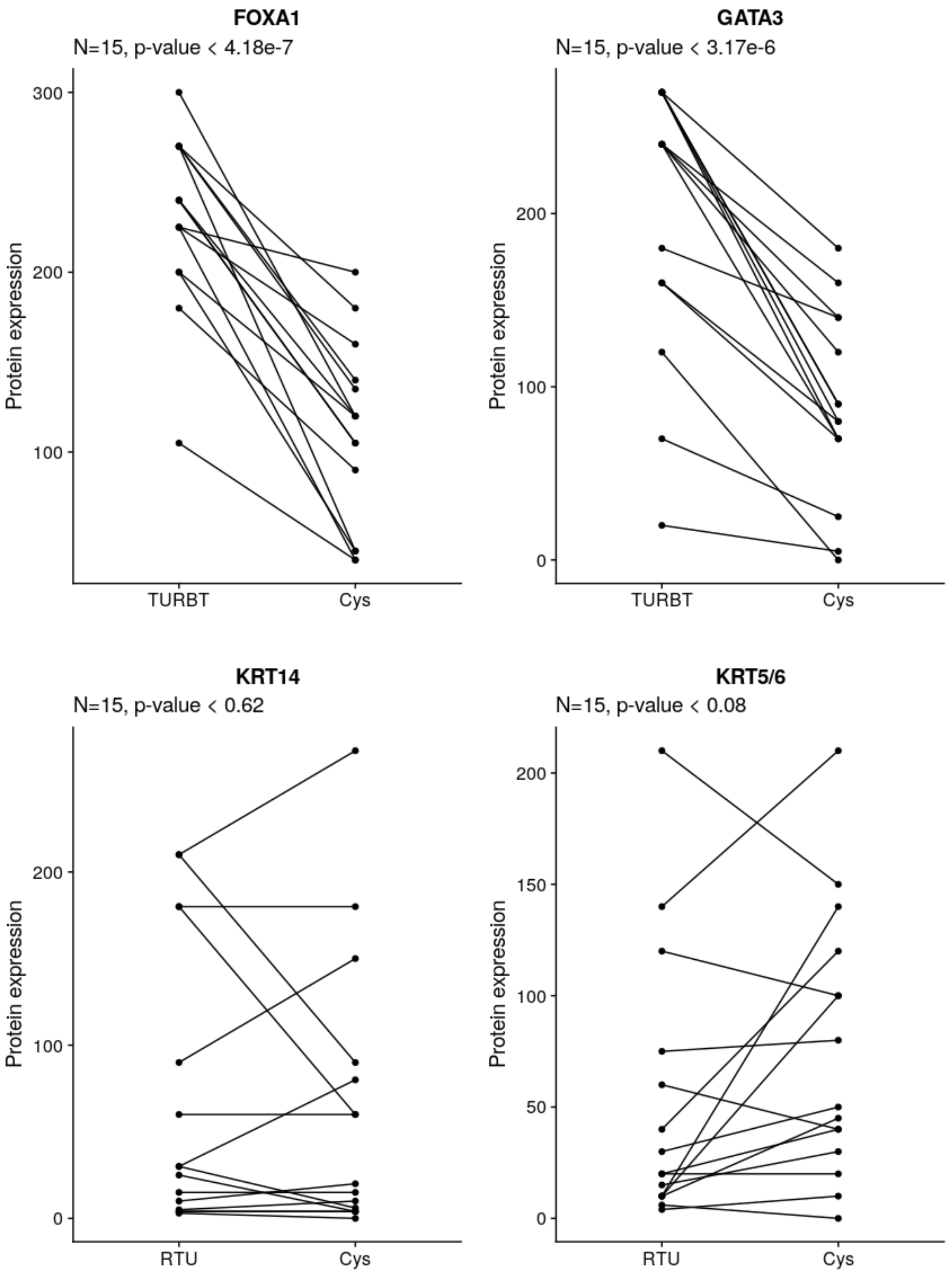

**Supplementary Figure 2**. Comparison of marker expression in full sections of samples obtained at TURBT and cystectomy (Cys) from patients who did not receive NAC. Statistical analysis: repeated measurements ANOVA.

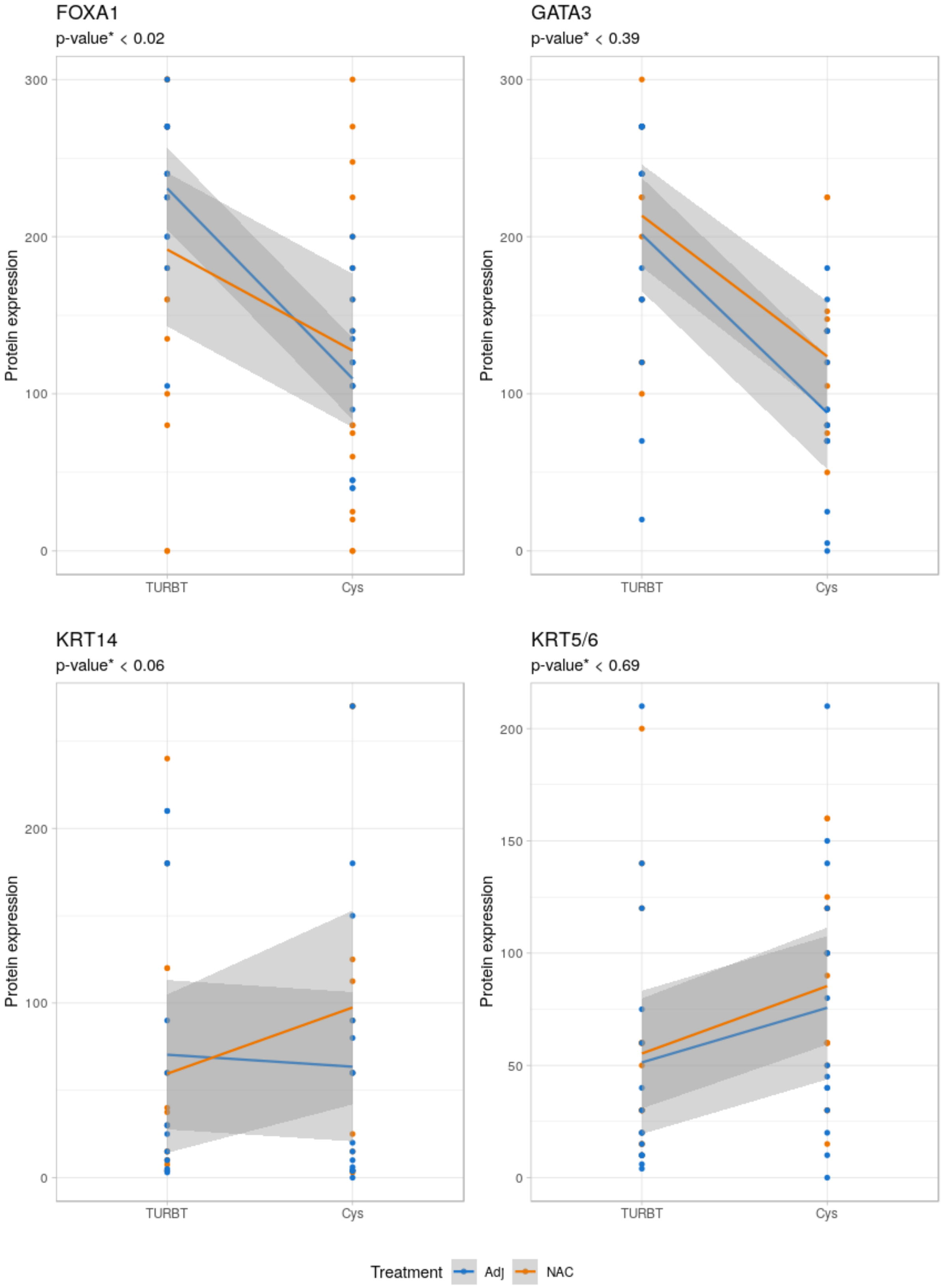

**Supplementary Figure 3**. Interaction analysis of marker expression in full sections of samples obtained at TURBT and cystectomy (Cys) from patients treated (orange) or not (blue) with NAC. Statistical analysis: linear mixed effects model; shown is the interaction P-value.
