## Supplementary figures and images for "Immunohistochemistry-based taxonomical classification of bladder cancer predicts response to neoadjuvant chemotherapy"

### Supplementary Figures

Supplementary Figure 1

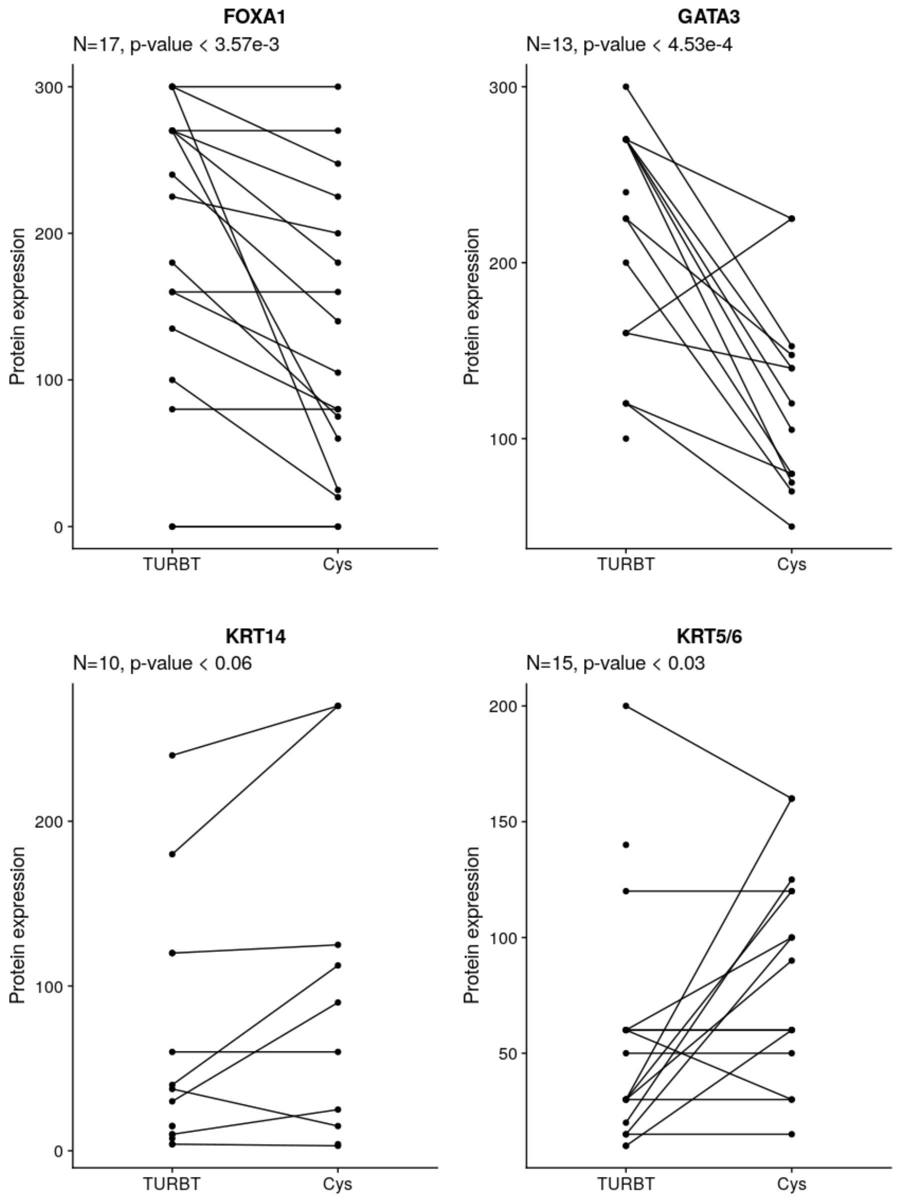

#### Supplementary Figure 2

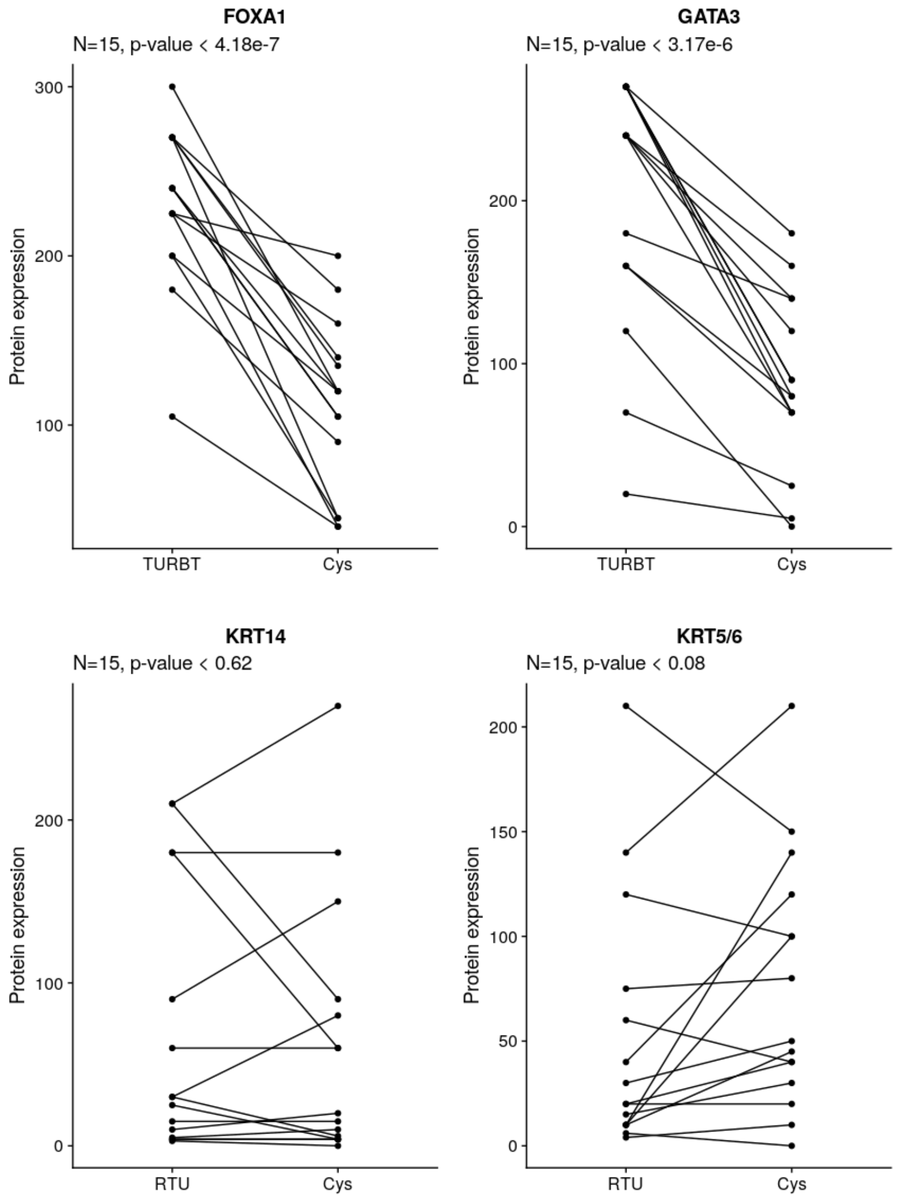

Supplementary Figure 3

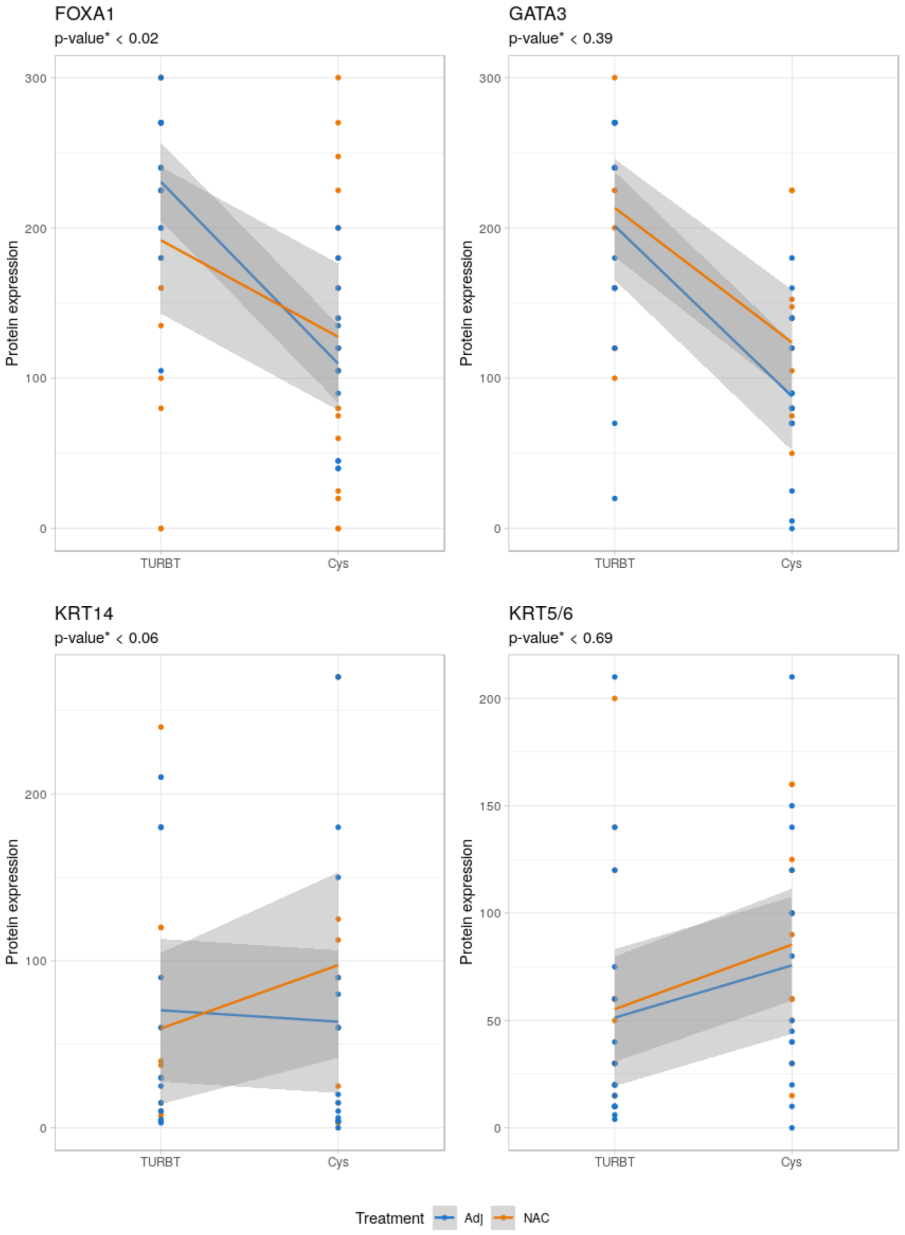
